# Targeted stimulation of motor cortex neural ensembles drives learned movements

**DOI:** 10.1101/2025.01.06.631504

**Authors:** Qiyu Chen, An Wu, Bin Yu, Soyoung Chae, Zijing Tan, Assaf Ramot, Johnatan Aljadeff, Takaki Komiyama

**Affiliations:** Department of Neurobiology, University of California San Diego, La Jolla, CA, USA; Center for Neural Circuits and Behavior, University of California San Diego, La Jolla, CA, USA; Department of Neurosciences, University of California San Diego, La Jolla, CA, USA; Halıcıoğlu Data Science Institute, University of California San Diego, La Jolla, CA, USA; Kavli Institute for Brain and Mind, University of California San Diego, La Jolla, CA, USA

## Abstract

During the execution of learned motor skills, the neural population in the layer 2/3 (L2/3) of the primary motor cortex (M1) expresses a reproducible spatiotemporal activity pattern. It is debated whether M1 actively participates in generating this activity pattern and whether this learned pattern causally drives the learned movement. To address these questions, we utilized *in vivo* two-photon calcium imaging combined with holographic optogenetic stimulation of functionally defined M1 L2/3 neuronal ensembles in mice performing a skilled lever-pressing task. A brief and synchronous stimulation of ∼20 neurons whose activity onset in voluntary trials precedes movement onsets induced movements that resembled the learned movement, while producing spatiotemporal activity patterns in other M1 neurons that resembled those during the voluntary learned movement. Moreover, trial-by-trial variability of optogenetically triggered population activity correlated with the variability in the induced movements. These trial-by-trial variabilities were predicted by the initial state of M1 population activity immediately preceding the stimulation. In some trials, the stimulation induced the learned activity without inducing overt movements, indicating that the learned activity is not simply a reflection of movements and instead can be induced internally. The stimulation failed to generate movements or learned activity when mice were disengaged from the task. Stimulation of the neurons whose activity followed voluntary movement onsets failed to induce the learned movement in task-engaged mice. Taken together, the learned activity pattern in M1 L2/3 can be generated when the M1 network is prepared at the optimal initial state and receives precise triggering inputs, supporting the active role of M1 in the generation of learned activity and learned movements.

## Main Text

Learning and generating motor skills are core functions of the brain, enabling animals to engage with their environment in a purposeful and adaptive manner. The primary motor cortex (M1) is a critical structure for these functions ^1–3^. Motor learning induces multiple forms of plasticity in M1, particularly in L2/3 neurons, which undergo extensive synaptic reorganization ^4–7^. This synaptic plasticity is accompanied by the emergence of a reproducible spatiotemporal pattern in the population activity of M1 L2/3 neurons during the execution of the learned motor skill ^4^.

Numerous studies have investigated the role of the learned activity pattern in M1 through neural recordings and loss-of-function experiments ^8–11^. However, it remains an open question how the learned activity pattern is generated. Furthermore, whether the learned activity pattern causally instructs the learned movement pattern is unknown. We addressed these issues by combining *in vivo* two-photon imaging with two-photon holographic optogenetic stimulation of movement-related neuronal ensembles in M1 L2/3 in mice performing a forelimb motor skill.

## Results

### Holographic stimulation of early-onset neuronal ensemble drives learned movement

We used a cued lever-press task, in which water-restricted mice were trained to grasp a lever and, in response to a sound cue delivered after a variable inter-trial interval, push and hold the lever for one second to acquire a water reward (**Fig. 1a**). After two weeks of daily training, mice became experts at this task, collecting rewards in most trials with short latencies (**Fig. 1b, c**).

**Fig. 1.**
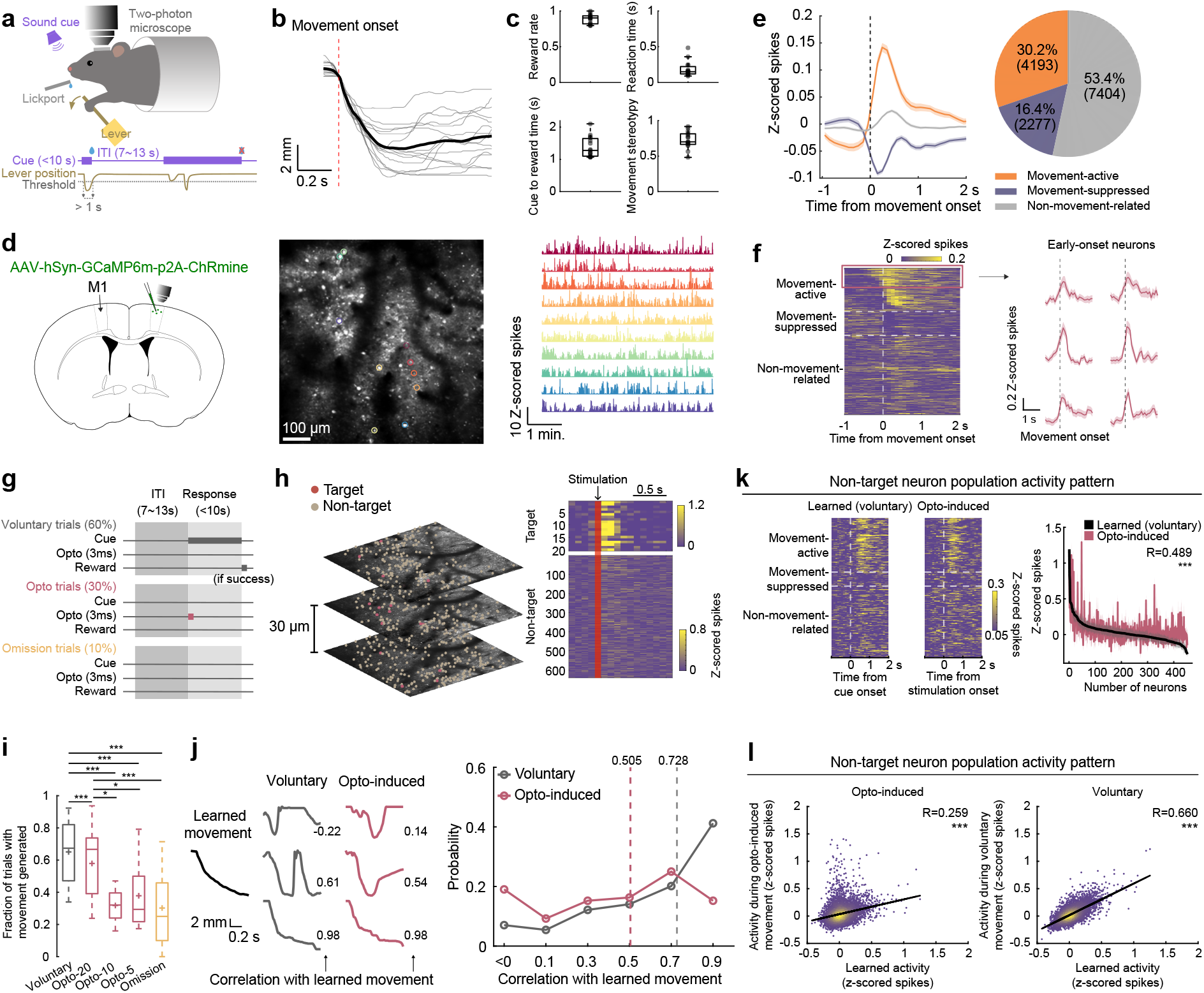
Holographic stimulation of early-onset neuronal ensemble drives learned movement. a. Schematic of experimental setup and task structure. ITI: inter-trial interval. b. Example lever trajectories of rewarded trials in an expert session from one mouse. Grey, individual trials; black, mean. c. Reward rate, reaction time, cue onset to reward time, and movement stereotypy (median trial-to-trial correlation coefficients of lever movements for each session) at expert stage (N = 15 mice). d. Left, schematic of viral injection into M1 to label L2/3 neurons. Middle, example field of view of *in vivo* two-photon calcium imaging. Right, z-scored time series of estimated spikes from example neurons outlined in the image in the middle. e. Left, average population activity of all movement-active, movement-suppressed and non-movement-related neurons. Mean ± SEM across session-averages. Right, proportions of movement-active, movement-suppressed and non-movement-related neurons. N = 13874 neurons from 25 sessions from 15 mice. f. Left, activity of movement-active (top, sorted by activity onset), movement-suppressed (middle), and non-movement-related (bottom) neurons averaged across movements, and aligned to movement onset. Vertical dashed line indicates movement onset. Each row represents one neuron. N = 13874 neurons from 25 sessions from 15 mice. Right, activity of example early-onset movement-active neurons averaged across movements, aligned to movement onset. Mean ± SEM. g. Trial structures for holographic stimulation during behavioral sessions. h. Left, example two-photon calcium imaging of three planes in M1 L2/3 spanning 60 μm in depth, showing target neurons (red) and non-target neurons (beige). Right, trial-averaged responses of early-onset target and non-target neurons in opto trials aligned to stimulation onset. Red vertical bar indicates stimulation timing. i. Fraction of trials with a movement generated within 0.3 sec from trial onset. Opto-20, 10, and 5 correspond to opto trials in which 20, 10, and 5 early-onset neurons were stimulated respectively (voluntary vs opto-20: *p* = 8.39×10^-5^, voluntary vs opto-10: *p* = 9.73×10^-7^, voluntary vs opto-5: *p* = 6.47×10^-6^, voluntary vs omission: *p* = 6.09×10^-26^, opto-20 vs opto-10: *p* = 0.010, opto-20 vs opto-5: *p* = 0.042, opto-20 vs omission: *p* = 1.23×10^-12^, opto-10 vs opto-5: *p* = 0.574, opto-10 vs omission: *p* = 0.698, opto-5 vs omission: *p* = 0.773, mixed-effects model). Box and whisker plots: median and interquartile range, + indicates mean. N = 25 sessions from 14 mice. j. Left, example movements from one session. Left, the learned movement; right, voluntary movements and opto-induced movements of various correlations with the learned movement. Right, probability of movements with varying correlations with the learned movement in voluntary and opto trials, respectively. Dashed lines indicate the medians (voluntary: 0.728, opto: 0.505). N = 15 sessions from 11 mice. k. Left, activity of movement-active, movement-suppressed, and non-movement-related neurons (separated by horizontal dashed lines) averaged across learned and opto-induced movements separately, from an example session. Each row represents a non-target neuron. Right, average activity of non-target neurons during the 1 sec following cue/stimulation onset sorted by amplitudes in learned movements from the same example session (Pearson’s correlation; *p*=1.54×10^-28^, R=0.489). l. Left, average activity of non-target neurons during voluntary learned movements versus during opto-induced movements pooled across sessions (*p* = 9.93×10^-135^, mixed-effects model; Pearson’s correlation, R=0.259). Each dot represents the activity of one neuron averaged during the 1 sec following cue/stimulation onset, averaged across trials. Right, average activity of non-target neurons during voluntary learned movements versus during all voluntary movements pooled across sessions (*p* < 1.00×10^-308^, mixed-effects model; Pearson’s correlation, R=0.660). Each dot represents the activity of one neuron averaged during the 1 sec following cue onset, averaged across trials. N = 15 sessions from 11 mice. For this and all other figures, *: *p*<0.05; **: *p*<0.01; ***: *p*<0.001.

Furthermore, the movement patterns became consistent across trials, a hallmark of learned motor skills (**Fig. 1c**). These features of motor learning are consistent with previous reports using similar tasks ^4–7,11–13^. It has been shown that the execution of the motor skill at this expert stage requires M1 activity ^11^. It was also found that training with these tasks over additional months can make the behavior independent of M1 ^11,14^ and so in the current study we focused our experiments on the early expert stage (2-3 weeks).

To record and stimulate the M1 L2/3 population, we injected a bicistronic AAV vector to co-express the calcium indicator GCaMP6m and the excitatory opsin ChRmine in the same neurons ^15^. We applied *in vivo* two-photon calcium imaging to record M1 L2/3 activity in task-performing mice (**Fig. 1d**). This approach revealed heterogeneous activity, including neurons showing significantly elevated activity (‘movement-active’, 30.2%) or decreased activity (‘movement-suppressed’, 16.4%) during movements (**Fig. 1e**). The activity onset of individual movement-active neurons relative to movement onset tiled the duration of movements. Of these neurons, we identified a group of ‘early-onset neurons’ (12.1 ± 1.5% of all neurons per session) whose activity onset preceded the movement onset (**Fig. 1f**).

We hypothesized that activity in these early-onset neurons contributed to the initiation of the learned movement. To test this hypothesis, we leveraged two-photon holographic optogenetic stimulation with a spatial light modulator that enables the activation of arbitrary groups of neurons within the field of view with single-cell resolution ^15–19^. We used this approach to ask whether an artificial stimulation of the early-onset neurons could induce the learned movement. We first performed two-photon calcium imaging for ∼60 trials, imaging at 3 depths within M1 L2/3 semi-simultaneously, each spaced 30 μm apart, to capture 555 ± 42 neurons. The data were quickly analyzed offline to identify early-onset neurons, after which the same imaging fields were identified, and the behavioral session was resumed. From this point forward, a majority of the trials were normal voluntary trials in which a lever press after the sound cue induced a water reward. In 30% of randomly interspersed trials (‘opto trials’), the sound cue was omitted, and instead a brief 3 ms pulse of optogenetic stimulation was delivered to either 5, 10, or 20 early-onset neurons simultaneously at the trial onset (see Methods). Critically, in opto trials, no reward was given even when mice made a successful movement. We also included omission trials, in which the sound cue and optogenetic stimulation were omitted, and no reward was ever given (**Fig. 1g**). In the opto trials, the stimulation successfully elicited reliable activity in the target neurons as revealed by simultaneous calcium imaging (**Fig. 1h**).

We observed that the stimulation of early-onset neurons can trigger movements. We focused our analysis on lever movements initiated within a short latency (0.3 sec) after trial onset (see Methods). Even though the fraction of trials in which movements were initiated was the highest in voluntary trials, fractions in 20-neuron opto trials were substantially higher than in omission trials. This indicates that movements in 20-neuron opto trials were indeed driven by the stimulation as opposed to voluntary movements serendipitously occurring after the stimulation (**Fig. 1i**). In contrast, 5- and 10-neuron opto trials did not induce movements more frequently than in omission trials, indicating that these were not sufficient stimulation to reliably induce movements. Consequently, we focused the following experiments and analysis on the 20-neuron paradigm. We next examined whether the opto-induced movements resembled the learned movement, which was defined for each session by identifying the subset of the voluntary movements that exhibited high stereotypy and averaging the movements in a random half of those trials. The trials used to construct the learned movement were excluded from subsequent analysis to avoid self-comparisons (see Methods). Both voluntary movements and opto-induced movements in individual trials showed variable correlations with the learned movement, with voluntary movements showing an overall higher correlation. Notably, however, about half of the opto-induced movements showed a high (>0.5) correlation with the learned movement (**Fig. 1j**). Thus, a brief and synchronous stimulation of ∼20 early-onset neurons frequently induced movements, many but not all of which resembled the learned movement.

How could 20 neurons drive the learned movement? M1 L2/3 is a highly recurrent network, where excitatory neurons form dense interconnections ^20–23^. We hypothesized that the activation of these ∼20 neurons elicit population activity in the other, non-target neurons that resembles the activity during the voluntary learned movement. To investigate this, we analyzed the trial-averaged activity of the non-target neurons during the voluntary trials in which the animal generated the learned movement (‘learned activity’) and compared that to the trial-averaged activity during opto-induced movements. We found that the activity of non-target neurons during opto-induced movements correlated with those during the voluntary learned movements (**Fig. 1k, l)**. In other words, neurons that were activated and suppressed during the voluntary learned movements tended to be also activated and suppressed during opto-induced movements, respectively. To explore the upper bound of this correlation, we repeated this analysis using the activity during voluntary trials (note that not all voluntary trials resulted in the learned movement, **Fig. 1j**). The voluntary trial activity correlated with the learned activity, similarly to (but more strongly than) the opto-induced activity (**Fig. 1l**). Thus, our stimulation of ∼20 early-onset neurons induced population activity patterns that resembled the learned pattern in the non-target neurons that are not directly stimulated.

### Stimulation of late-onset neuronal ensemble fails to drive learned movement

It is striking that our artificial stimulation of a small group of neurons induced naturalistic population activity. One possibility is that any random input can engage the local pattern-completion network and induce the same naturalistic activity to drive movements. To test this, we conducted another set of experiments, this time targeting a subset of ‘late-onset neurons’ (15.9 ± 0.9% of all neurons per session), whose activity onsets were after movement onset in voluntary movements (**Fig. 2a, b**). ∼20 of the late-onset neurons were randomly selected as target neurons, matching the number stimulated for the early-target stimulation experiments. The stimulation was done during behavioral sessions in which a majority of trials were voluntary trials, in the same way as early-target stimulation sessions.

**Fig. 2.**
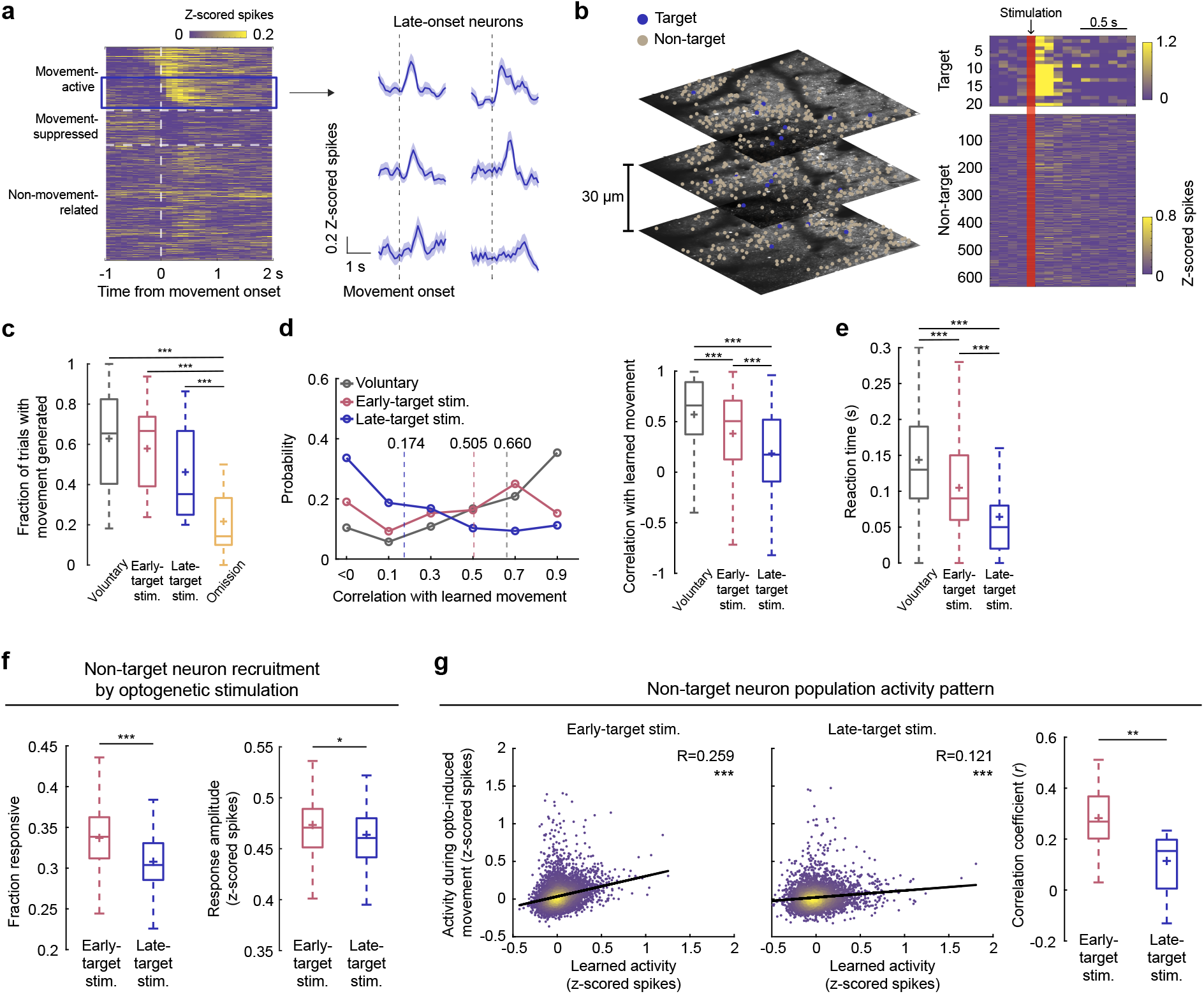
Stimulation of late-onset neuronal ensemble fails to drive learned movement. a. Left, activity of movement-active (top, sorted by activity onset), movement-suppressed (middle), and non-movement-related (bottom) neurons averaged across movements, and aligned to movement onset. Vertical dashed line indicates movement onset. Each row represents one neuron. N = 13874 neurons from 25 sessions from 15 mice. Right, activity of example late-onset movement-active neurons averaged across movements, aligned to movement onset. b. Left, example two-photon calcium imaging of three planes in M1 L2/3 spanning 60 μm in depth, showing target neurons (blue) and non-target neurons (beige). Right, trial-averaged responses of late-onset target and non-target neurons in opto trials aligned to stimulation onset. Red vertical bar indicates stimulation timing. c. Fraction of trials with a movement generated within 0.3 sec from trial onset (voluntary vs early: *p* = 4.85×10^-5^, voluntary vs late: *p* = 0.095, voluntary vs omission: *p* = 6.93×10^-35^, early vs late: *p* = 0.179, early vs omission: *p* = 7.93×10^-18^, late vs omission: *p* = 1.93×10^-7^, mixed-effects model). d. Left, probability of movements with varying correlations with the learned movement for each trial type. Dashed lines indicate the medians (voluntary: 0.660, early: 0.505, late: 0.174). Right, correlation with the learned movement in each trial type (voluntary vs early: *p* = 5.24×10^-12^, voluntary vs late: *p* = 1.23×10^-12^, early vs late: *p* = 2.45×10^-4^, mixed-effects model). Box and whisker plots: median and interquartile range, + indicates mean. e. Reaction time of movements in each trial type (voluntary vs early: *p* = 1.10×10^-11^, voluntary vs late: *p* = 1.55×10^-30^, early vs late: *p* = 1.98×10^-6^, mixed-effects model). Box and whisker plots: median and interquartile range, + indicates mean. f. Left, fraction of responsive non-target neurons per trial for early-target and late-target stimulation trials (*p* = 5.82×10^-5^, mixed-effects model). Right, response amplitudes of the responsive non-target neurons for early-target and late-target stimulation trials (*p* = 0.018, mixed-effects model). g. Left, average activity of non-target neurons during voluntary learned movements versus in early-target stimulation trials pooled across sessions (*p* = 9.93×10^-135^, mixed-effects model; Pearson’s correlation, R=0.259). Each dot represents the activity of one neuron averaged during the 1 sec following cue/stimulation onset, averaged across trials. Middle, average activity of non-target neurons during voluntary learned movements versus in late-target stimulation trials pooled across sessions (*p* = 7.86×10^-20^, mixed-effects model; Pearson’s correlation, R=0.121). Each dot represents the activity of one neuron averaged during the 1 sec following cue/stimulation onset, averaged across trials. Right, correlation coefficients between non-target neuron population activity in voluntary learned movements and opto-induced movements calculated for individual sessions for early-target and late-target stimulation sessions, respectively (*p* = 2.1×10^-3^, Wilcoxon rank-sum test). For this figure, N = 15 early-target stimulation sessions from 11 mice and 10 late-target stimulation sessions from 8 mice with the exception of **Fig. 2a**.

Late-target stimulation induced movements in a subset of stimulation trials, showing a non-significant trend toward lower efficacy compared to voluntary and early-target trials (**Fig. 2c**). We next examined the quality of stimulation-induced movements by evaluating the correlation with the learned movement. We found that the movements induced by late-target stimulation were significantly less similar to the learned pattern than the movements in voluntary and early-target stimulation trials (**Fig. 2d**). Thus, compared to early-target stimulation, late-target stimulation did not reliably induce the learned movement. These results indicate that the choice of target neurons is important—early-target stimulation is much more effective at inducing the learned movement than late-target stimulation.

We also observed differences in the reaction time of movement initiation across trial types. Specifically, opto-induced movements in both early-target and late-target stimulation trials had a significantly shorter reaction time than voluntary movements (**Fig. 2e**). The shorter reaction time supports the idea that holographic stimulation of M1 L2/3 neurons bypassed the sensory-motor transformation taking place in voluntary trials, leading to faster movement onsets. Interestingly, the reaction time was shorter in late-target stimulation than in early-target stimulation. This raises the possibility that early-target stimulation induced an additional circuit computation to perform pattern completion to generate the learned activity and movement.

We next explored the neural underpinnings that differentiate behavioral outcomes between early-target and late-target stimulation. To this end, we analyzed the activity of the non-target neurons within M1 L2/3 and how they were recruited by the stimulation of the target neurons. We found that early-target stimulation led to the activation of a larger fraction of non-target neurons, and higher response amplitudes of the recruited non-target neurons compared to late-target stimulation (**Fig. 2f**). Furthermore, we found that the population activity induced by late-target stimulation did not resemble the learned activity as much as early-target stimulation (**Fig. 2g**).

Overall, stimulation of size-matched late-onset ensembles failed to reliably induce the learned movement, likely because the elicited population activity patterns did not sufficiently match the activity pattern underlying the learned movement. These findings suggest that generating the learned movement requires the activation of early-onset neuronal ensembles as the triggering input to the M1 network, which then propagate the activity to other nearby neurons to complete the learned activity pattern.

Given the distinct behavioral and neural outcomes between early-target and late-target stimulation, we next asked whether these effects could be attributed to confounding differences in the properties of the target neurons. M1 neurons exhibit distinct input-output connectivity, somatotopic organization and functional specification depending on their anatomical location. To account for this, we assessed the depth, anterior-posterior, and medial-lateral distributions of early and late targets, and confirmed that they were not significantly different from each other (**Extended Data Fig. 1a**). The degree of spatial clustering among neurons may reflect distinct local circuit organizations, where stimulation of a clustered ensemble could be more effective at generating functional output ^24^. However, we found that early and late targets exhibited comparable degrees of spatial clustering which were similar to the overall M1 L2/3 population revealed by shuffling labels (**Extended Data Fig. 1b**). Other than their activity onset timing that differs by definition, their activity properties during voluntary movements were similar, as quantified by the peak amplitude and response consistency (**Extended Data Fig. 1c**).

Collectively, these other properties of early- and late-onset target neurons are unlikely to explain the distinct outcomes of their stimulation.

### M1 activity explains movement variability

Opto-induced movements exhibited varying degrees of similarity to the learned movement pattern across trials. We next examined whether this variability is reflected in the variability of induced population activity. We focused on the early-target stimulation sessions where we were successful at triggering movements that resembled the learned movement in a considerable fraction of trials. We applied principal component analysis (PCA) on the non-target population activity (voluntary trials only) and then projected the population activity of individual trials onto the neural space that captured 80% of the variance (see Methods). In voluntary trials, on a trial-by-trial basis, there was a negative correlation between the population activity distance to the learned activity and movement correlation with the learned movement (**Fig. 3a**). In other words, the more similar the population activity was to the learned activity, the more similar the movement was to the learned movement. A similar result was observed in opto trials, where movements with low correlations with the learned movement showed population activity trajectories distinct from the learned activity (**Fig. 3b**). These results were not affected when we included the target neurons in the analysis (**Extended Data Fig. 2**). These findings suggest that the learned movement was generated when the induced population activity resembled the learned activity.

**Fig. 3.**
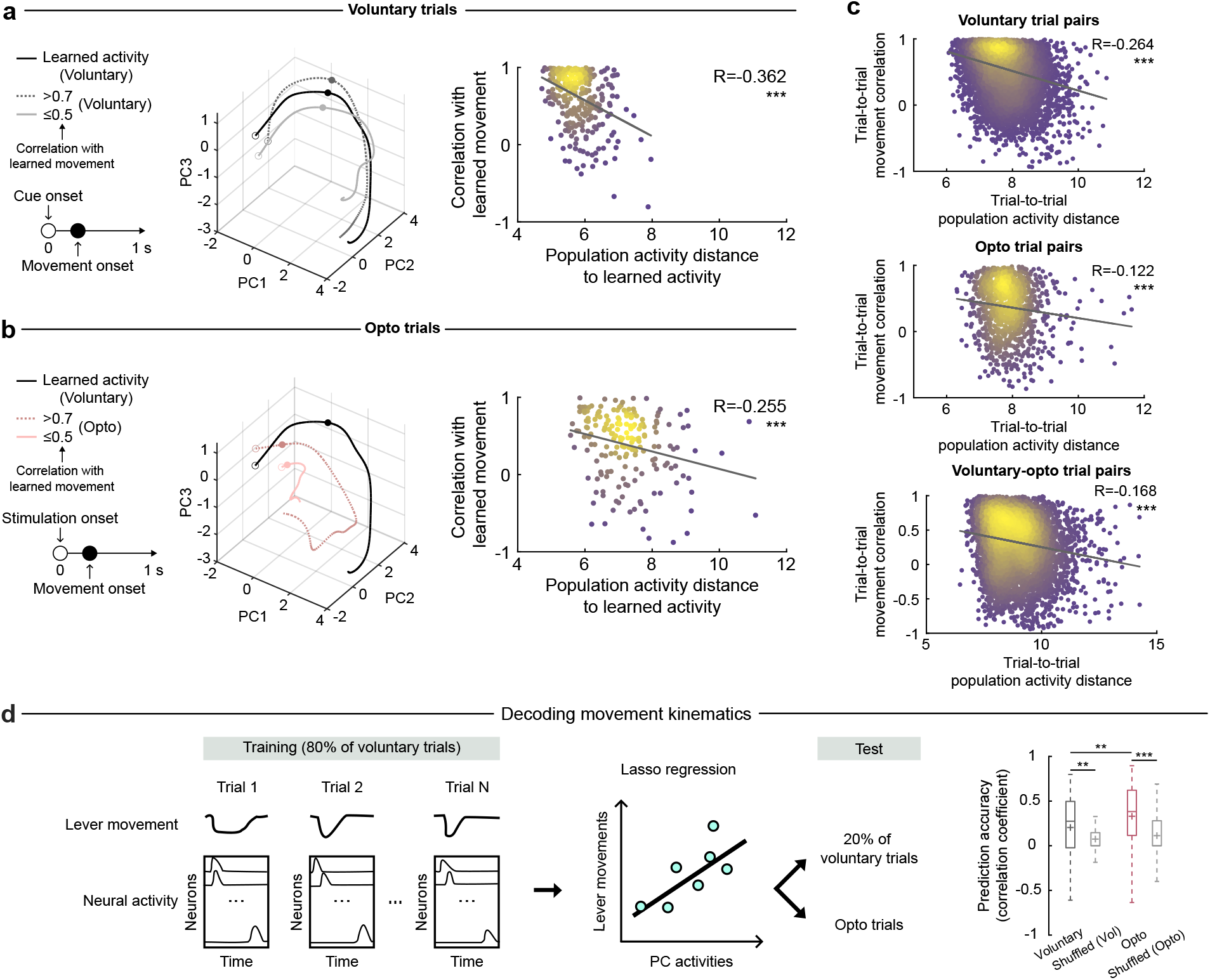
M1 activity explains movement variability. **a.** Left, population activity trajectories of all non-target neurons in voluntary trials of an example session, for the 1 second after cue onset. Black, learned activity; grey dotted and solid lines, trials with high (>0.7) and low (≤0.5) correlation with the learned movement respectively. Right, population activity distance to the learned activity negatively correlated with correlation with the learned movement (*p* = 3.32×10^-11^, mixed-effects model; Pearson’s correlation, R=-0.362). Each dot represents an individual trial. **b.** Left, population activity trajectories of all non-target neurons in opto trials of an example session, for the 1 second after stimulation onset. Black, learned activity; pink dotted and solid lines, trials with high (>0.7) and low (≤0.5) correlation to the learned movement respectively. Right, population activity distance to the learned activity negatively correlated with correlation with the learned movement (*p* = 4.53×10^-4^, mixed-effects model; Pearson’s correlation, R=-0.255). Each dot represents an individual trial. **c.** Population activity distance negatively correlated with movement correlation between voluntary trials (top, *p* = 5.90×10^-90^, mixed-effects model; Pearson’s correlation, R = - 0.264), between opto trials (middle, *p* = 5.63×10^-5^, mixed-effects model; Pearson’s correlation, R = -0.122), and between voluntary and opto trials (bottom, *p* = 1.77×10^-24^, mixed-effects model; Pearson’s correlation, R = -0.168). **d.** Left, schematic illustrating the prediction of lever movements on individual trials based on M1 population activity. A regression model was trained using 80% of voluntary trials and tested on the remaining voluntary trials as well as opto trials (see Methods). Right, prediction accuracy, measured by correlation coefficients between the actual and predicted lever movements, were significantly better than chance in both voluntary and opto trials (voluntary vs shuffled: *p* = 5.38×10^-3^, opto vs shuffled: *p* = 5.84×10^-13^, mixed-effects model), and better in opto trials compared to voluntary trials (*p* = 1.39×10^-3^, mixed-effects model). Box and whisker plots: median (voluntary: 0.275, voluntary shuffled: 3.56×10^-17^, opto: 0.383, opto shuffled: 6.41×10^-17^) and interquartile range, + indicates mean. For this figure, N = 15 sessions from 11 mice.

In addition to the similarity to the learned movement, a previous study showed that trial-by-trial variability in movement kinematics is represented in M1 L2/3 activity of trained animals ^4^. Thus, we next examined the activity-movement relationship in pairs of trials to ask whether the relationship is shared across both the voluntary and opto-induced movements. Specifically, we examined voluntary trial pairs, opto trial pairs, and voluntary-opto trial pairs to assess whether the movement similarity and the population activity similarity correlate with each other across voluntary and opto-induced movements. In all three comparisons, we observed a significant negative correlation between the population activity distance and the movement correlation (**Fig. 3c**). This suggests that M1 L2/3 activity reflects the movement kinematics on a trial-by-trial basis, regardless of whether the movement was generated voluntarily or by our optogenetic stimulation.

We further explored the activity-movement relationship using a decoding analysis to decode the movement time series on individual trials with population activity using lasso regression. The decoder was trained using 80% of voluntary trials and tested on the other trials (**Fig. 3d left**, see Methods). The decoder was able to decode the voluntary movements on the held-out trials substantially better than the chance level defined by trial shuffles (**Fig. 3d right**). This confirms that M1 L2/3 activity contains information about movement kinematics. When the same decoder trained on voluntary trials was tested on opto trials, we again saw that it was able to decode movements substantially better than chance (**Fig. 3d right**). This suggests that the mechanism by which M1 L2/3 drives movements was shared between voluntary and opto trials. Surprisingly, the decoder performance was significantly better in opto trials than in held-out voluntary trials, even though the decoder was trained with voluntary trials. This may suggest that voluntary movements are also influenced by additional sources of variability outside of M1 which dilutes the coupling between M1 activity and movement, while opto movements are more purely controlled by M1 activity.

Taken together, these findings indicate that the variability in M1 L2/3 activity can explain the variability of both voluntary and opto-induced movements.

### Initial state of M1 activity predicts movement variability

What accounts for the variability in the non-target population activity in opto trials, when the input was the same in every trial (stimulation of the same ∼20 early-onset neurons)? Previous studies have found that the variability in the state of motor cortex population activity prior to movement initiation contributes to movement variability ^25–29^. Thus, we hypothesized that the initial population state before the stimulation influences the subsequent evolution of population activity, thereby contributing to movement variability. To address this hypothesis, we examined the population activity during the period (0.5-sec) immediately prior to our stimulation in opto trials or cue onset in voluntary trials. In opto trials, we observed that the pre-stimulation population activity that was closer to the pre-cue state of the learned activity was associated with subsequent, opto-induced movements more similar to the learned movement (**Fig. 4a**). In other words, the pre-stimulation activity state predicted whether the subsequent stimulation was going to be successful in inducing the learned movement. Similar results were found in voluntary trials, such that the pre-cue population state predicted the quality of the voluntary movements (**Fig. 4b**). These results were consistent whether or not the target neurons were included in the analysis (**Extended Data Fig. 2**). Thus, in both the opto and voluntary trials, on a trial-by-trial basis, there appears to be an optimal initial state that facilitated the subsequent generation of the learned movement. This is consistent with the ‘initial condition hypothesis’ which posits that the initial state of a dynamical system partially defines the evolution of the subsequent spatiotemporal activity ^29^. Supporting this notion, the pre-stimulation and pre-cue activity state predicted the subsequent population activity during movements on a trial-by-trial basis, in both the opto and voluntary trials respectively (**Fig. 4c**). Perhaps as a consequence, we found a negative correlation between the distance of initial states and movement correlation—in pairs of voluntary trials, pairs of opto trials, and pairs of the two trial types (**Fig. 4d**).

**Fig. 4.**
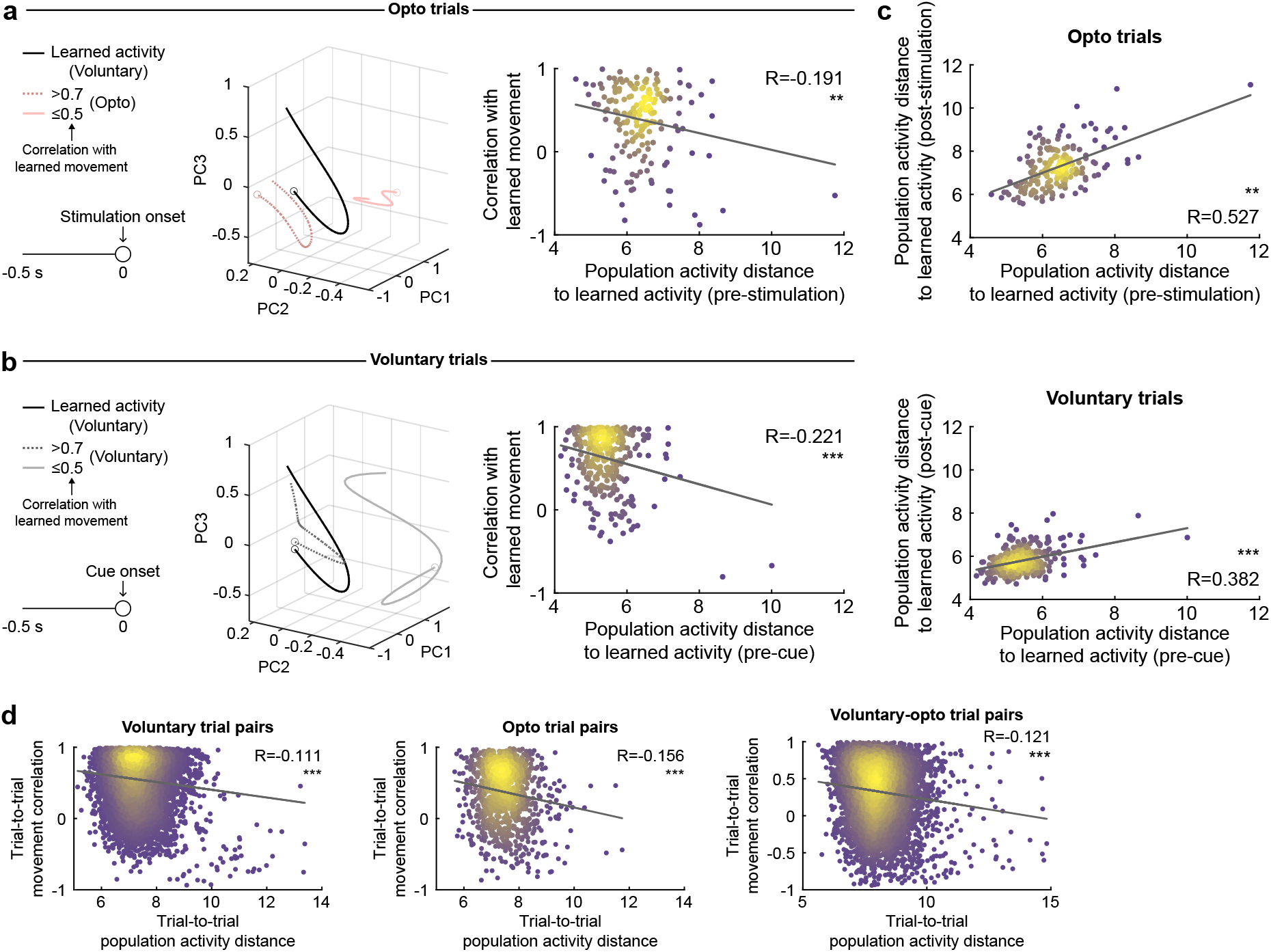
Initial state of M1 activity predicts movement variability. a. Left, population activity trajectories of all non-target neurons in opto trials of an example session, for the 0.5 second period before stimulation onset. Black, learned activity; pink dotted and solid lines, trials with high (>0.7) and low (≤0.5) correlation to the learned movement respectively. Right, population activity distance to the learned activity negatively correlated with correlation with the learned movement (*p* = 0.009, mixed-effects model; Pearson’s correlation, R = -0.191). Each dot represents an individual trial. b. Left, population activity trajectories of all non-target neurons in voluntary trials of an example session, for the 0.5 second period before cue onset. Black, learned activity; grey dotted and solid lines, trials with high (>0.7) and low (≤0.5) correlation with the learned movement respectively. Right, population activity distance to the learned activity negatively correlated with correlation with the learned movement (*p* = 7.68×10^-5^, mixed-effects model; Pearson’s correlation, R = -0.221). Each dot represents an individual trial. c. Population activity distance to the learned activity showed a correlation between pre- and post-stimulation periods in opto trials (top, *p* = 0.002, mixed-effects model; Pearson’s correlation, R = 0.527) and between pre- and post-cue periods in voluntary trials (bottom, *p* = 1.08×10^-8^, mixed-effects model; Pearson’s correlation, R = 0.382). d. Population activity distance before the cue or stimulation negatively correlated with correlation of subsequent movements between voluntary trials (left, *p* = 7.65×10^-14^, mixed-effects model; Pearson’s correlation, R = -0.111), between opto trials (middle, *p* = 3.00×10^-4^, mixed-effects model; Pearson’s correlation, R = -0.156), and between voluntary and opto trials (right, *p* = 1.16×10^-13^, mixed-effects model; Pearson’s correlation, R = -0.121). For this figure, N = 15 sessions from 11 mice.

Taken together, the initial state of the M1 L2/3 population predicts the variability in subsequent M1 activity and movements.

### Optogenetically triggered population activity is not a consequence of movements

When optogenetic stimulation was delivered to ∼20 early-onset neurons, we observed activity in the non-target M1 L2/3 population that correlated with movement variability. This could be because the activity of target neurons propagates to other L2/3 neurons to generate the population activity. However, an alternative possibility is that the target neurons directly engage downstream circuits to generate movements, largely bypassing the local L2/3 network. In this scenario, M1 L2/3 population activity may reflect sensory feedback signals related to the resulting movement. To parse these possibilities, we leveraged the subset of trials in which early- or late-target stimulation failed to evoke any overt movements (**Fig. 2c**), which provided a window into network dynamics in the absence of motor output.

For both early- and late-target stimulation, the non-target neurons exhibited substantial activity in no-movement trials, indicating that stimulation of 20 neurons can generate population activity in non-target neurons without engaging movements, even though this activity tended to be less prolonged than in movement trials (**Extended Data Fig. 3**). We analyzed this short-latency (0.3 sec) population activity both in terms of the activity amplitude and correlation with the learned activity pattern. In early-target stimulation, both the amplitude and correlation of population activity were similar in movement and no-movement trials (**Fig. 5a**). Comparing non-target population activity between early-versus late-target stimulation, we found that early-target stimulation evoked stronger activity that better resembled the learned activity than late-target stimulation in both movement and no-movement trials (**Fig. 5b**). These observations indicate that the distinct neural responses in early-versus late-target stimulation arise from their differences in how they engage the local circuit, instead of as a consequence of the differences in the induced movements. Furthermore, they suggest that the learned activity induced by early target stimulation is not purely a consequence of induced movements (e.g. sensory feedback) and rather completed internally in the brain.

**Fig. 5.**
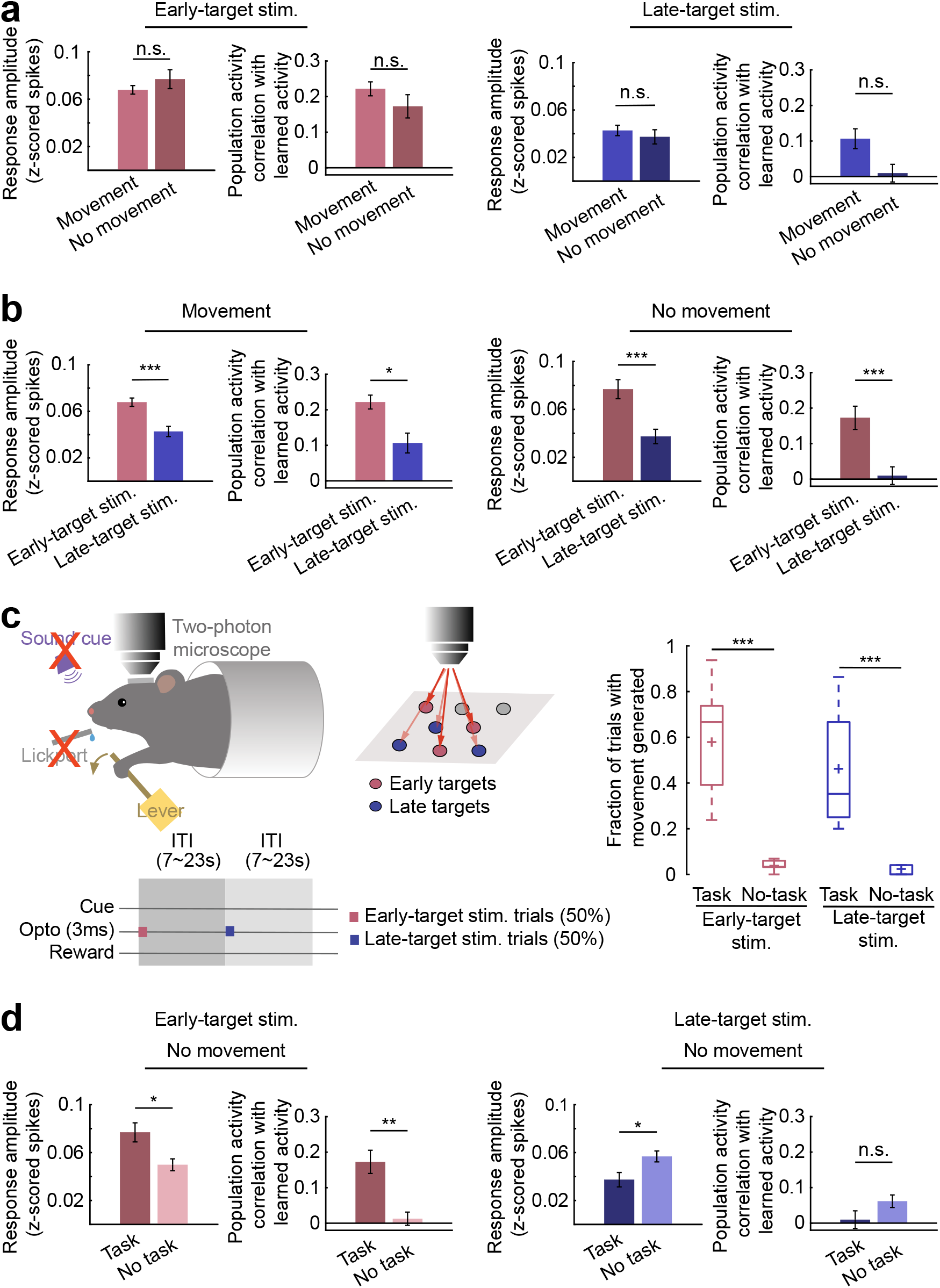
Early target stimulation can induce learned activity without inducing movements in a context-dependent manner. a. Left, activity amplitude and correlation with learned activity of non-target neurons during movement trials versus no-movement trials following early-target stimulation (amplitude: *p* = 0.274, correlation: *p* = 0.251, mixed-effects model). Activity of neurons was averaged during the 0.3 sec following stimulation onset, averaged across trials. Right, same as left for late-target stimulation (amplitude: *p* = 0.464, correlation: *p* = 0.800, mixed-effects model). b. Left, activity amplitude and correlation with learned activity of non-target neurons during movement trials following early-versus late-target stimulation (amplitude: *p* = 1.27×10^-5^, correlation: *p* = 0.031, mixed-effects model). Right, same as left for no-movement trials (amplitude: *p* = 3.05×10^-4^, correlation: *p* = 2.92×10^-4^, mixed-effects model). c. Left, schematic of no-task condition setup and task structure. Right, fraction of trials with movements generated within 0.3 sec from trial onset (early task vs early no-task: p = 1.48×10^-11^, late task vs late no-task: p = 6.68×10^-6^, mixed-effects model). Box and whisker plots: median and interquartile range, + indicates mean. d. Left, activity amplitude and correlation with learned activity of non-target neurons during no-movement trials following early-target stimulation under task versus no-task condition (amplitude: *p* = 0.045, correlation: *p* = 0.008, mixed-effects model). Right, same as left for late-target stimulation (amplitude: *p* = 0.027, correlation: *p* = 0.112, mixed-effects model).

### Optogenetic induction of learned movement requires task engagement

We have shown that optogenetic stimulation of early-onset targets can induce the learned activity pattern and learned movement. Further, generation of learned activity and movement depended on the initial state of M1. All the experiments so far were done in task-engaged animals that were presumably preparing the learned movement. We asked whether this behavioral context is important for the observed effects of our stimulation. To address this, we performed stimulation in a no-task condition in which well-trained mice were presented with the lever but received no sound cue and had no access to the lickport. We then delivered the same optogenetic stimulation used in the task context—interleaving early-target and late-target stimulation—each followed by a variable interval (**Fig. 5c**). In this no-task context, both early- and late-target stimulation failed to evoke any movement reliably (**Fig. 5c**), demonstrating that task engagement is critical for the stimulation to produce motor outputs. We then examined non-target population activity following stimulation, comparing task and no-task contexts specifically in no-movement trials.

Task engagement bi-directionally modulated the non-target response: early-target stimulation produced stronger activity in the task context, whereas late-target stimulation elicited weaker activity compared to the no-task condition. Moreover, the population activity evoked by stimulation in the no-task context did not resemble the learned activity pattern (**Fig. 5d**). Taken together, these findings suggest that stimulation of early-onset neurons drives movements in a context-dependent manner, in which task engagement primes the network to selectively express the learned activity pattern in response to inputs to the early-onset neurons.

## Discussion

In this study, we leveraged holographic stimulation to gain insights into the mechanisms that regulate the learned activity in M1 L2/3 and its role in generating learned movements. We found that brief and synchronous stimulation of approximately 20 early-onset, but not late-onset, movement-related neurons often triggered the learned activity within M1 L2/3 and induced the learned movement in a task-dependent manner. The efficacy of this stimulation depended on the initial state of the M1 L2/3 population activity. Taken together, we propose that the execution of the learned movement is ensured by two conditions: first, the motor cortex prepares the movement by entering the appropriate initial state of the population activity. Second, M1 receives specific inputs from other brain area(s) that activate the early-onset neurons. When both of these two conditions are met, M1 generates the learned activity pattern that reliably drives the learned movement.

The importance of the initial state aligns well with the dynamical systems perspective on motor cortex movement control ^28,30,31^. Previous studies have found that the pre-movement activity state of motor cortex correlates with movement parameters ^26,29,30^, leading to the postulate that the motor cortex functions as a dynamical machine in which its state at one point in time predicts the next state. Our results provide a direct support for this idea by demonstrating that the network response to our identical stimulus is variable depending on the initial state of the network.

However, the initial state does not completely specify the subsequent activity, as the external inputs also affect the evolution of network activity shown by the differences observed in our early-target vs. late-target stimulation. Future studies could explore whether targeted perturbations can correct suboptimal initial states to enhance movement consistency.

M1 exhibits dynamic activity patterns during movements. Whether M1 plays an active role in generating the activity pattern or passively reflects input activity has been debated ^28,32–34^. By directly stimulating a subset of task-relevant neurons in M1 L2/3, we demonstrated that locally initiated cortical activity can ‘pattern-complete’ to drive the learned activity pattern, even in the absence of overt movements. We argue that these observations exclude the possibility that M1 is a purely passive machine that only reflects the temporally dynamic activity pattern provided by inputs from other brain areas. One possibility is that learning shapes the recurrent connectivity within M1 L2/3, forming a local circuit that can autonomously generate the learned activity pattern. In this possibility, activity may initiate in a group of neurons and then propagates through the network like a chain reaction—reminiscent of models proposed in zebra finch song production ^35,36^. Furthermore, the inputs from the motor thalamus likely provide the triggering signal, activating the early-onset ‘starter cells’, and this thalamocortical connectivity refines during learning ^36–41^. Computational studies have shown that recurrent neural networks are capable of generating complex and initial-condition-dependent activity patterns that underlie movements ^42–45^. M1 L2/3, as the primary input layer of M1 ^46^, is highly recurrent in its connectivity and shows a high degree of learning-related plasticity ^4–7, 20–23^, further supporting this possibility. Future research should elucidate the relative contributions of local vs. global interactions in cortical pattern generation and how particular circuits can be selectively primed in a task-dependent manner.

## Acknowledgments

We cherish our memories of our dear friend and colleague, An Wu, and dedicate this manuscript to her surviving parents, Mr. Wu and Ms. Qu. We thank Bobbie Morales, Ashley Medina, David Arakelyan, and Elanore Hall for technical assistance; and other Komiyama Lab members, especially J. Li and E. Gjoni, for discussion. We also thank all our friends who have reached out to us to offer support during the difficult time.

## Author contributions

Conceptualization: AW, TK

Methodology: AW, QC, TK, JA

Investigation: QC, AW, BY, ZT, AR

Visualization: QC

Formal analysis: QC, AW, SC

Validation: QC

Resources: AW, QC, TK

Software: QC, AW

Data curation: QC

Funding acquisition: TK

Project administration: TK

Supervision: TK

Writing – original draft: QC, TK

Writing – review & editing: QC, TK, JA, BY, SC, ZT, AR

