## Supplemental Information for "Targeted stimulation of motor cortex neural ensembles drives learned movements"

#### Methods

##### Animals

All procedures were conducted in compliance with protocols approved by the UCSD Institutional Animal Care and Use Committee (IACUC) and guidelines of the National Institutes of Health (NIH). C57BL/6 mice were acquired from Charles River Laboratory. Animals were group-housed in disposable plastic cages with standard bedding in a room with a reversed light cycle. Experiments were performed during the dark period (10:00-22:00). Males and females were randomly used for experiments. The experiments were done on young adults (8 – 15 weeks old).

##### Surgery and virus injections

Mice were anesthetized with 3% isoflurane for induction, maintained at 1% throughout the surgical procedure. To prevent infection and inflammation, Baytril (10 mg/kg) and dexamethasone (2 mg/kg) were administered subcutaneously. A custom-built headplate was secured to the exposed skull using superglue and dental cement. A craniotomy (~3 mm in diameter) was performed over the right caudal forelimb area of M1, centered at 0.3 mm anterior and 1.5 mm lateral from the bregma. Viral vectors (AAV8-hSyn-GCaMP6m-p2A-ChRmine-Kv2.1-WPRE, Addgene) were diluted to $1.6\times10^{12}$ vg/mL and injected via a glass pipette at a depth of ~250 μm from the pia. Multiple injections (3-4) of 20–30 nl were made. Following injections, a glass window was implanted into the craniotomy and affixed with Vetbond (3M), superglue, and dental cement. At the end of the surgery, buprenorphine (0.1 mg/kg) was administered subcutaneously for postoperative pain management.

##### Behavior

Water restriction began ~4 weeks post-surgery and ~15 days prior to behavioral training. The water volume was gradually reduced until stabilizing at 1 mL per day. Animal weights were monitored daily to ensure maintenance of at least 75% of their pre-restriction body weight.

Mice were trained to perform a lever-press task under head-fixation one session per day. The lever consisted of a rotary encoder (E6B2-CWZ3E, YUMO) attached to a brass rod, which was held in place by two opposing springs and could be moved horizontally. The lever position was continuously recorded at 50k Hz using WaveSurfer and at 100 Hz using Bpod (Sanworks), integrated with custom codes to monitor threshold crossings. Bpod also controlled the behavioral program. Mice grasped the lever with their left paw while resting their right paw on a stationary thick rod, with their bodies in a plexiglass tube. A tone (2 kHz) signaled the cue period (up to 10 s), during which a successful lever press was rewarded with water (~5 μl per trial). Each trial was followed by an inter-trial interval (ITI) of variable duration (7-13 s). A successful lever press was defined as the lever crossing a set threshold (0.83~1.25 mm) and remaining beyond the threshold for at least 1 second. Failure to achieve a rewarded movement during the cue period was not punished. Lever presses during ITI were neither rewarded nor punished. Each training session consisted of ~100 trials.

During the no-task condition (**Fig. 5c**), mice were first trained with the task for 2 weeks. These mice were then placed in the behavioral setup with the lever accessible, but the lickport was removed and no sound cue nor reward was provided. Optogenetic stimulation was applied to early targets and late targets in an interleaved manner, with each stimulation followed by a variable inter-trial interval (7-23 sec).

##### *In vivo* two-photon imaging

Imaging was conducted with a commercial two-photon microscope (Ultima 2Pplus, Bruker) equipped with a 16×/0.8 NA objective (Nikon) and a Ti-Sa laser (MaiTai, Newport) tuned to 925 nm. Image acquisition was controlled by Prairie View Software (Bruker). Images (512 × 512 pixels at ×1.5 zoom) were recorded at ~10 Hz per plane across three planes, spaced 30 μm apart in depth. The microscope was equipped with an electrotunable lens (Bruker) to enable imaging at different z-planes. Frame times were recorded and synchronized with behavioral recordings using the WaveSurfer software.

##### Holographic stimulation with a spatial light modulator (SLM) during behavior

A pre-opto session was conducted while mice performed the lever-press task under two-photon imaging. The pre-opto session included approximately half the number of trials in a regular training session to prevent the animals from reaching satiety before the opto session. Mice were then returned to their home cage while the neuronal activity was analyzed offline to assess functional properties and determine which neurons to target in the subsequent opto sessions. After the analysis, mice were put back under the microscope, the same imaging fields were identified, and target neurons were manually selected for stimulation. Typically, the interval between the pre-opto and opto sessions was less than an hour. For early-target stimulation sessions, we sorted movement-active neurons by activity onset timing and selected from the earliest neurons that were visible and healthy. Similarly, for late-target stimulation sessions, we prioritized the neurons with later activity onsets. The activity onsets of target neurons were -9 ± 13 ms and 238 ± 24 ms (mean ± SD) around movement onset in early-target and late-target stimulation sessions, respectively. We selected ~20 neurons for stimulation based on the observation that stimulating fewer than 20 neurons rarely elicited movements (**Fig. 1i**). Additionally, in most experiments, we did not have many more than 20 early-onset neurons that we were able to visually identify quickly while defining the spatial patterns for stimulation.

For holographic stimulation, the microscope and acquisition system (Bruker) was used with the NeuraLight 3D Spatial Light Modulator (SLM) (Bruker) and a second, 1035-nm high power laser (Monaco 1035-40-40, Coherent). Spatial accuracy of SLM targeting was ensured through built-in calibration routines prior to each stimulation session. The stimulation laser operated at a pulse repetition rate of 1 MHz and spirally scanned over each target cell body (10 μm diameter, 5 revolutions), delivering ~7 mW of power per cell for 3 ms in each opto trial.

#### Data analysis

##### Behavior analysis

Lever position was recorded by both Bpod and WaveSurfer. WaveSurfer recordings were used whenever available. Lever position was sampled at 100 Hz. For movement analysis, we excluded trials that started during ongoing movements, defined as lever speeds exceeding 12.5 mm/s during the 50 ms prior to trial onset. For trials with no ongoing movements, movement onset was defined as the first time point after trial onset where lever velocity exceeded 41.7 mm/s in the forward direction. Reaction time was calculated as the interval from trial onset to movement onset. Trials without a movement onset within 0.3 seconds of trial onset were excluded from further analysis. We imposed this constraint because movements occurring much later than the stimulation in opto trials were unlikely to be triggered by the stimulation. The specific 0.3-second time window was chosen based on the mean reaction time of 0.24 seconds observed in expert mice during voluntary movements (**Fig. 1c**). For consistency, the same criterion was applied to movements in all trial types.

For analyses of no-movement trials (**Fig. 5**), we applied stricter criteria to ensure the absence of overt movements. The threshold for movement onset was defined as the first time point after trial onset at which the lever velocity exceeded 25.0 mm/s in the forward direction. Trials in which no such movement onset occurred within 1-second following stimulation were included in these analyses as no-movement trials.

Movement analysis was done on lever time series for 1-sec starting from 0.1 s before movement onset. The learned movement pattern was defined for each session separately to account for the idiosyncrasies of individual animal behavior. We first identified highly stereotyped movements as those with a correlation above 0.7 with the mean of all voluntary movements. A random half of these stereotyped movements were averaged to form the learned movement. For comparisons involving the learned movement, the trials used in their construction were excluded from the analysis.

#### Image analysis

*Regions of interest (ROI) identification and fluorescence analysis*

Suite2P software [47] was used for motion correction and regions of interest (ROI) extraction, which correspond to individual neurons. ROI identification was further refined by manual selection. Fluorescence signals from ROIs were deconvolved to remove fluorescence decay and infer spiking activity using Suite2P built-in deconvolution algorithm. The estimated spike activity was used for all neural activity analyses.

In order to match neurons between pre-opto sessions and opto sessions, ROIMatchPub (https://github.com/ransona/ROIMatchPub) was used for automatic detection, followed by manual matching of target neurons with custom code.

*Movement-related neuron classification*

Movement-related neurons were classified as previously described [4,6,12]. Briefly, the dot product of the binarized lever movement time series (movement epochs versus non-movement epochs) and estimated spikes was calculated for each neuron. This value from actual data was then compared to the dot products when shuffling the time points of movement epochs 1000 times. Actual values above the 95th percentile of the shuffled distribution were classified as ‘movement-active’, while those below the 5th percentile were classified as ‘movement-suppressed’. All other neurons were considered as ‘non-movement-related’.

*Movement-related activity onset*

To estimate the activity onset timing of individual neurons during movements, we aligned the activity of each neuron to movement onsets. We only considered movements that did not have another movement within 1 s before movement onset to avoid contamination of activity related to the other movements. Activity averaged across movements was then used for onset detection. Activity onset was defined as the first frame between -0.2 s before movement onset to end of movement in which activity was higher than baseline activity (average activity between -1 s to -0.5 s before movement onset) by more than one standard deviation for 200 ms.

*Non-target neuron population activity pattern*

Activity of non-target neurons were averaged during the 1-second period following cue or stimulation onset. Time-averaged activity of each neuron was then averaged across learned movements, opto-induced movements, or voluntary movements (**Fig. 1k, l and Fig. 2g**).

*Non-target neuron recruitment*

Response amplitudes of non-target neurons to optogenetic stimulation of target neurons were calculated for each non-target neuron in each trial as the difference between the average activity during the 1-second period following stimulation onset and the average activity during the 1-second period preceding stimulation onset. Non-target neurons with response amplitudes exceeding 0.2 z-scored spikes were classified as responsive to, and thus recruited by, the optogenetic stimulation of the target neurons (**Fig. 2f**).

*Target neuron general properties*

The anatomical distribution of targets was quantified for each session as the fraction of targets along the imaging depth, anterior–posterior, and medial–lateral axes, and these values were subsequently averaged across sessions. The degree of spatial clustering was quantified by computing, for each target neuron, the distance to its nearest neighbor target neuron. The resulting nearest-neighbor distance distribution was then compared to the chance distribution generated by shuffling target labels 1,000 times. Peak amplitudes of movement-related activity were quantified as the maximum neural activity within the −1 to 2 s window around movement onset. Response consistency was defined as the fraction of movements in which a given neuron's activity within the 2-s window following movement onset was significantly higher than its baseline activity. For both peak amplitudes and response consistency measurements, we used movements recorded during the pre-opto session in the absence of any stimulation (**Extended Data Fig. 1**).

*Principal component analysis (PCA)*

For each session, we compiled a data matrix for voluntary trials, $X$, of size $n \times fr \times tr$, where $n$ is the number of non-target neurons, $fr$ is the 40 image frames corresponding to -2 s to 2 s around cue onset, and $tr$ is the total number of voluntary trials. PCA was applied to trial-averaged data in $X$ (size $n \times fr$) and the principal components (PCs) explaining 80% of the variance in neural space were extracted. We then projected population activity of individual trials to the PCs explaining 80% of total variance (**Fig. 3-5, Extended Data Fig. 2**).

*Movement-activity relationship*

To calculate the trajectory distance, we computed the Euclidean distance in the PC space between trajectories at each time point and these values were added across time points. For pre-cue/pre-stimulation activity, we averaged the distance over the 500 ms preceding trial onset, while for post-cue/post-stimulation activity, we averaged the distance during the 1 second following trial onset to obtain the trajectory distance for each trial. To construct the learned trajectory, we averaged the projected activity from the trials that contributed to the learned movement. The activity from these trials was then excluded from further analyses to avoid self-comparison (**Fig. 3-4, Extended Data Fig.2**).

*No-movement trial analysis*

Response amplitudes were calculated as the trial-averaged activity of non-target neurons averaged during the 0.3-second period following stimulation onset. To calculate correlation with learned activity, we computed the Pearson correlation between the projected population activity averaged over the 0.3-s window following stimulation onset in opto trials and that of the learned movement (**Fig. 5**).

**Movement decoding analysis**

To examine the relationship between movement-related activity and movements, we sought to decode across-trial variability of lever movements using across-trial variability of population activity.

We aligned the temporal resolution of lever traces with that of the neural activity by downsampling lever traces from 100 Hz to 10 Hz by averaging every 10 data points. We divided voluntary trials into 80% for training of the model and 20% for testing. Opto trials were not included in training.

To define variability in lever movements, we utilized the difference of the lever movement on each trial from the learned movement. Similarly, for population activity, we utilized the difference of the trial activity from the learned activity. These differences from the learned patterns, or 'residuals', were used instead of the original activity and movement to avoid building a model that only predicts the trial-averaged movement and does not explain across-trial variabilities.

The population activity residuals were first processed by principal component analysis (PCA), concatenating activity across trials to capture trial-by-trial variability. The top principal components (PCs) capturing 80% of the total variance were selected. Using these PC activities in the training set, we built a LASSO regression model to predict lever residuals. The optimal lambda value was determined through 10-fold cross-validation. The trained model was tested on the test voluntary trials and opto trials. Model performance was evaluated by calculating correlation coefficients between actual and predicted lever residuals on a trial-by-trial basis.

To test statistical significance, a control regression model was built. Lever trace residuals were randomly shuffled across trials while preserving their temporal structure, while neural residuals retained their original trial order. The lever trace residuals were shuffled such that trials were never retained in their original positions. The control model was then trained and tested as above.

Finally, we compared the correlation coefficients from the main model with those obtained from the control model using a mixed-effects model, accounting for session effects. Additionally, model performance for test voluntary trials and opto trials was evaluated within the same mixed-effects framework.

#### Statistics

All statistical analyses were performed in MATLAB. Statistical tests used are described in figure legends. Statistical significance was determined by a standard threshold of p-values less than 0.05. Results are denoted in figures as follows: $p < 0.05$ as *, $p < 0.01$ as **, and $p < 0.001$ as ***.

For analyses pooling trials across sessions, we used mixed-effects models (using the fitlme function in MATLAB) to account for the nested data structure and minimize the effects of session-to-session variability. The models used in this manuscript are as follows:

$$y \sim trial\ type + (1|mouse{:}session)$$

where the fixed effect is the trial type, voluntary or early-target stimulation or late-target stimulation or omission trials, movement or no-movement trials, task trials or no-task trials, and the random effect is experimental session nested within each mouse. This model was used to compare the fraction of trials with movement generated (**Fig 1i, 2c, Fig. 5c**), correlation with learned movement (**Fig. 2d**), reaction time (**Fig. 2e**), fraction of responsive non-target neurons (**Fig. 2f**), response amplitude of non-target neurons (**Fig. 2f**), population activity amplitudes (**Fig. 5a-b, d**), population activity correlation with learned activity (**Fig. 5a-b, d**), activity peak amplitudes and response consistency during movement (**Extended Data Fig. 1c**), between pairs of trial types.

$$y \sim learned\ activity + (1|\ mouse{:}session)$$

where the fixed effect is the learned activity, and the random effect is the experimental session nested within each mouse. This model was used to assess relationship between learned activity and opto-induced or voluntary movement activity (**Fig. 1l, 2g**).

$$y \sim distance + (1|\ mouse{:}session)$$

where the fixed effect is the population activity distance, and the random effect is the experimental session nested within each mouse. This model was used to assess relationship between population activity distance to learned activity and correlation with learned movement (**Fig. 3a-b, 4a-b, Extended Data Fig. 2**), relationship between trial-to-trial population activity distance and trial-to-trial movement correlation (**Fig. 3c, 4d**), and the relationship between pre-stimulation/pre-cue population activity distance to learned activity and post-stimulation/post-cue population activity distance to learned activity, respectively (**Fig. 4c**).

$$y \sim condition + (1|\ mouse{:}session)$$

where the fixed effect is the condition, voluntary or shuffled (voluntary) or opto or shuffled (opto), and the random effect is the experimental session nested within each mouse. This model was used to compare the correlation coefficients between actual and predicted lever residuals from different regression models (**Fig. 3f**).

$$y \sim group*location + (1|mouse:session)$$

where the fixed effects were group, location, and their interaction (group × location), and the random effect was the experimental session nested within each mouse. The interaction term was used to evaluate whether the two groups exhibited comparable spatial distributions of target neurons across cortical depth (**Extended Data Fig. 1a, left**), anterior–posterior position (**Extended Data Fig. 1a, middle**), and medial–lateral position (**Extended Data Fig. 1a, right**).

$$y \sim data\ type + (1|mouse:session)$$

where the fixed effect was data type and the random effect was the experimental session nested within each mouse. This model was used to evaluate differences in spatial clustering between early-target neurons and shuffled data (**Extended Data Fig. 1b, left**), between late-target neurons and shuffled data (**Extended Data Fig. 1b, middle**), and between early- and late-target neurons (**Extended Data Fig. 1b, right**).

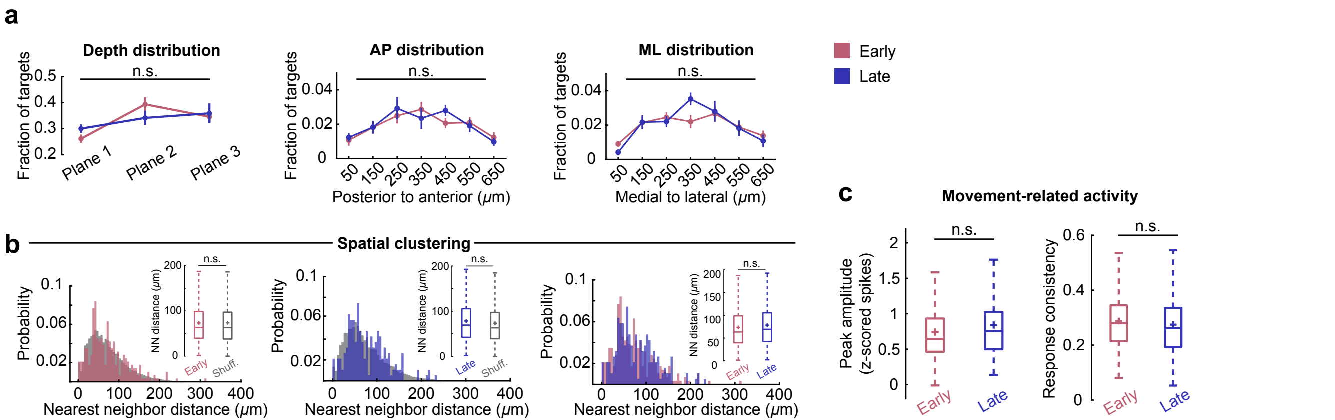


**Extended Data Fig. 1**

**Extended Data Fig. 1. General properties of target neurons do not explain their distinct stimulation outcomes**

a. Left, fraction of early- or late-onset targets located in different imaging depths ($p$ = 0.153, mixed-effects model). Middle, fraction of early- or late-onset targets located along the anterior-posterior axis within the field of view ($p$ = 0.667, mixed-effects model). Right, fraction of early- or late-onset targets located along the medial-lateral axis ($p$ = 0.813, mixed-effects model).

b. Left, the distribution of nearest neighbor distances between early-onset targets and a shuffled control ($p$ = 0.930, mixed-effects model). Middle, same as left for late-onset ($p$ = 0.133, mixed-effects model). Right, Comparison between early- and late-onset targets ($p$ = 0.262, mixed-effects model).

c. Left, the peak activity amplitude of early- vs. late-onset targets during voluntary movements ($p$ = 0.351, mixed-effects model). Right, the response consistency of early- vs. late-onset targets during voluntary movements ($p$ = 0.516, mixed-effects model). Response consistency was measured as the fraction of movements during which a given neuron was active.

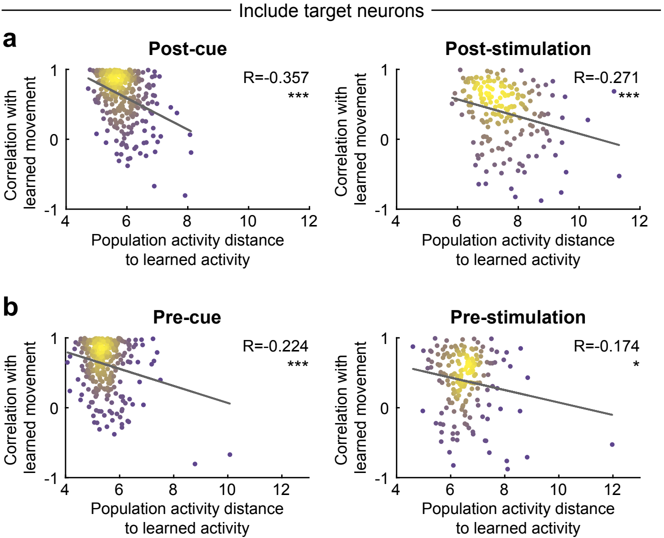


**Extended Data Fig. 2**

**Extended Data Fig. 2. Activity-movement relationship analysis including target neurons**

These are equivalent to **Fig. 3a, b** and **Fig. 4a, b** except that the analysis includes target neuron activity.

a. Left, post-cue population activity distance to the learned activity negatively correlated with correlation with the learned movement in voluntary trials ($p = 6.84\times10^{-11}$, mixed-effects model; Pearson's correlation, R = -0.357). Each dot represents an individual trial. Right, post-stimulation population activity distance to the learned activity negatively correlated with correlation with the learned movement in opto trials ($p = 1.79\times10^{-4}$, mixed-effects model; Pearson's correlation, R = -0.271).

b. Left, pre-cue population activity distance to the learned activity negatively correlated with correlation with the learned movement in voluntary trials ($p = 6.06\times10^{-5}$, mixed-effects model; Pearson's correlation, R = -0.224). Each dot represents an individual trial. Right, pre-stimulation population activity distance to the learned activity negatively correlated with correlation with the learned movement in opto trials ($p = 0.017$, mixed-effects model; Pearson's correlation, R = -0.174).

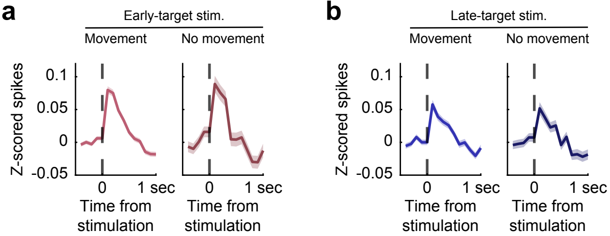


### Extended Data Fig. 3

**Extended Data Fig. 3. Population activity in non-target neurons in no-movement trials.**

a. Average population activity of non-target neurons in early-target stimulation movement trials (left) and no-movement trials (right). Mean ± SEM pooled across sessions.
b. Same as a but for late-target stimulation.
